# Crossmodal integration improves sensory detection thresholds in the ferret

**DOI:** 10.1101/014407

**Authors:** Karl J. Hollensteiner, Florian Pieper, Gerhard Engler, Peter König, Andreas K. Engel

## Abstract

During the last two decades ferrets (*Mustela putorius*) have been established as a highly efficient animal model in different fields in neuroscience. Here we asked whether ferrets integrate sensory information according to the same principles established for other species. Since only few methods and protocols are available for behaving ferrets we developed a head-free, body-restrained approach allowing a standardized stimulation position and the utilization of the ferret’s natural response behavior. We established a behavioral paradigm to test audiovisual integration in the ferret. Animals had to detect a brief auditory and/or visual stimulus presented either left or right from their midline. We first determined detection thresholds for auditory amplitude and visual contrast. In a second step, we combined both modalities and compared psychometric fits and the reaction times between all conditions. We employed Maximum Likelihood Estimation (MLE) to model bimodal psychometric curves and to investigate whether ferrets integrate modalities in an optimal manner. Furthermore, to test for a redundant signal effect we pooled the reaction times of all animals to calculate a race model. We observed that bimodal detection thresholds were reduced and reaction times were faster in the bimodal compared to unimodal conditions. The race model and MLE modeling showed that ferrets integrate modalities in a statistically optimal fashion. Taken together, the data indicate that principles of multisensory integration previously demonstrated in other species also apply to crossmodal processing in the ferret.

## Introduction

During the last two decades ferrets (*Mustela putorius*) have become increasingly relevant as an animal model in different fields in neuroscience [1–24]. Ferrets have been domesticated for over 2000 years and are easy to handle and train on behavioral tasks [15,25–29]. As a carnivore ferrets have excellent visual and auditory sensing and are well suited for cross-modal integration studies. An additional advantage is that the ferret brain shows substantial homologies with that of other animal models established in neuroscience, such as the cat [10,11,18–20] and the primate [26]. Extensive work has been performed to map cortical and subcortical regions of the ferret brain functionally and anatomically [3,11,17–20,22]. These mapping studies have shown that ferrets have highly complex sensory cortical systems, making them an interesting model for the study of sensory processing pathways, response properties and topographies of sensory neurons. Several studies have addressed multisensory response properties in anesthetized ferrets [2,4,8,14], but multisensory interactions have not yet been studied in a behavioral preparation in this species.

Substantial effort has been made to uncover principles of multisensory integration in a variety of species and paradigms [30–35]. Multisensory integration is crucial for animals and influences behavior in synergistic or competitive ways. Sensory integration can lead to faster reaction times, better detection rates and higher accuracy values in multi-compare to unimodal conditions [33,36,37]. Specifically, sensory integration increases the reliability by reducing the variance in the sensory estimate [36,38,39]. The consistent estimate with the lowest variance is the Maximum Likelihood Estimate (MLE) [40], which can be derived from the weighted sum of the individual sensory estimates, with weights being inversely proportional to the variance of the unisensory signals [36,39]. A substantial number of studies indicate that humans and animals indeed integrate information across sensory modalities in this way [33,36,38,39,41–46]. For example, Ernst and Banks [36] used a MLE model to predict the results of a visual-haptic experiment and showed that humans integrate information in a statistically optimal fashion. Similar results were obtained by application of MLE in a human audio-visual study [37] and in a vestibular-visual study in macaque monkeys [47]. These studies demonstrate that the MLE is a robust statistical model to predict the crossmodal response and to test whether subjects integrate information in a statistically optimal fashion. As a results of the sensory integration process, the accumulation of information in multimodal compared to unimodal conditions is faster, which in turn leads to decreased reaction times (RT) [48–53].

In the present study, we investigated whether ferrets integrate sensory signals according to the same principles established for humans [33,54] and non-human primates [47]. Previous studies in behaving ferrets have used either freely-moving [13,15,55] or head-restrained [26] animals. Here, we developed a head-free, body-restrained approach allowing a standardized stimulation position and the utilization of the ferret’s natural response behavior. An additional demand was that the setup should be sufficiently flexible to allow combination of the behavioral protocol with electrophysiological recordings. We established a behavioral paradigm, requiring combination and integration in the auditory and/or visual modality, to investigate features of uni- and multisensory integration in the ferret and compare it to data reported from other species. Ferrets were tested in a 2-alternative-choice task requiring them to detect lateralized auditory, visual, or combined audio-visual targets of varying intensity. We expected the ferrets to perform more accurate and faster in the bimodal cases, because congruent inputs from two modalities provide more reliable sensory evidence. We first determined unimodal thresholds for auditory amplitude and visual contrast detection. Subsequently, we combined both modalities and compared psychometric fits and the RTs between all conditions. We used MLE to model psychometric curves and to probe whether ferrets integrate visual and auditory signals in an optimal manner. Furthermore, to test for a redundant signal effect (RSE) we pooled the RT of all animals in order to calculate a race model and to investigate potential intensity- and modality-dependent effects [49,56,57].

## Materials and Methods

Ferrets were trained in a spatial detection paradigm, which was used to perform two behavioral experiments. In the first experiment, the animals’ auditory and visual unisensory detection thresholds were determined. In the second experiment, unimodal and bimodal thresholds were assessed in a combined approach, using the unimodal results from the first experiment to adjust the test parameters.

### Animals

Four adult female ferrets (*Mustela putorius;* Euroferret, Dybbølsgade, Denmark), aged 2 years (n=2) and 4 years (n=2), from two different litters were tested in the experiment. They were individually housed in a standard ferret cage with enriched environment under controlled ambient conditions (21°C, 12-h light/dark cycle, lights on at 8:00 a.m.). The animals had ad libidum access to food pellets. Access to tap water was restricted 8h before the experiments and the training procedure. All behavioral testing was done during the light cycle between 10:00 a.m. and 2:00 p.m.

### Ethics statement

All experiments were approved by the Hamburg state authority for animal welfare (BUG-Hamburg; Permit Number: 22/11) and performed in accordance with the guidelines of the German animal protection law. All sections of this report adhere to the ARRIVE Guidelines for reporting animal research [58].

### Experimental setup

The experiments were carried out in a dark sound attenuated chamber to ensure controlled conditions for sensory stimulation. Once per day each ferret performed the experimental task in a custom-build setup (Fig. 1A,D). We crafted a flat-bottomed tube to conveniently house the animal during the experiment. The body of the ferret was slightly restrained by fixation to three points in the tube via a harness, while the head remained freely movable outside the tube throughout the session. The semi-circular tube was fixed on an aluminum pedestal to level the animals’ head at 20cm distance to the center of the LED screen used for visual stimulation (BenQ XL2420T, Taipei, Taiwan). On the front (‘head side’), two convex aluminum semicircles were mounted horizontally below and above the animals’ head, respectively, at 150mm distance. They carried three light-barrier-fibers (FT-FM2), in the center, left and right, respectively, connected to high-speed (sampling interval: 250μs) receivers (FX301, SUNX, Aichi, Japan). This allowed the detection of the animal head during the experiments. In addition, a waterspout was co-localized with each light-barrier source. On both sides of the LED screen a speaker (T1; Beyerdynamic, Heilbronn, Germany) was placed with a 45° angle to the screen surface and at the height of the horizontal screen midline. A custom made 3-channel water-dispenser was installed outside the sound attenuated chamber to avoid acoustical interference during the experiments. It consisted of three valves from SMC Corporation (Tokyo, Japan), a Perfusor syringe (Melsungen, Germany) as water reservoir and Perfusor tubing to connect it with the waterspouts. The setup was controlled by custom-made routines using the Matlab environment (The Mathworks Inc.; MA, USA) on a MacPro. Behavioral control (light-barriers) and reward application (water-dispenser) were triggered through NI-PCI-cards (NI-6259 and NI-6251; National Instruments GmbH, Munich, Germany). The Psychtoolbox and the custom-written NI-mex-function referred to the same internal clock allowing the precise timing of behavior and stimulation .

**Figure 1.**
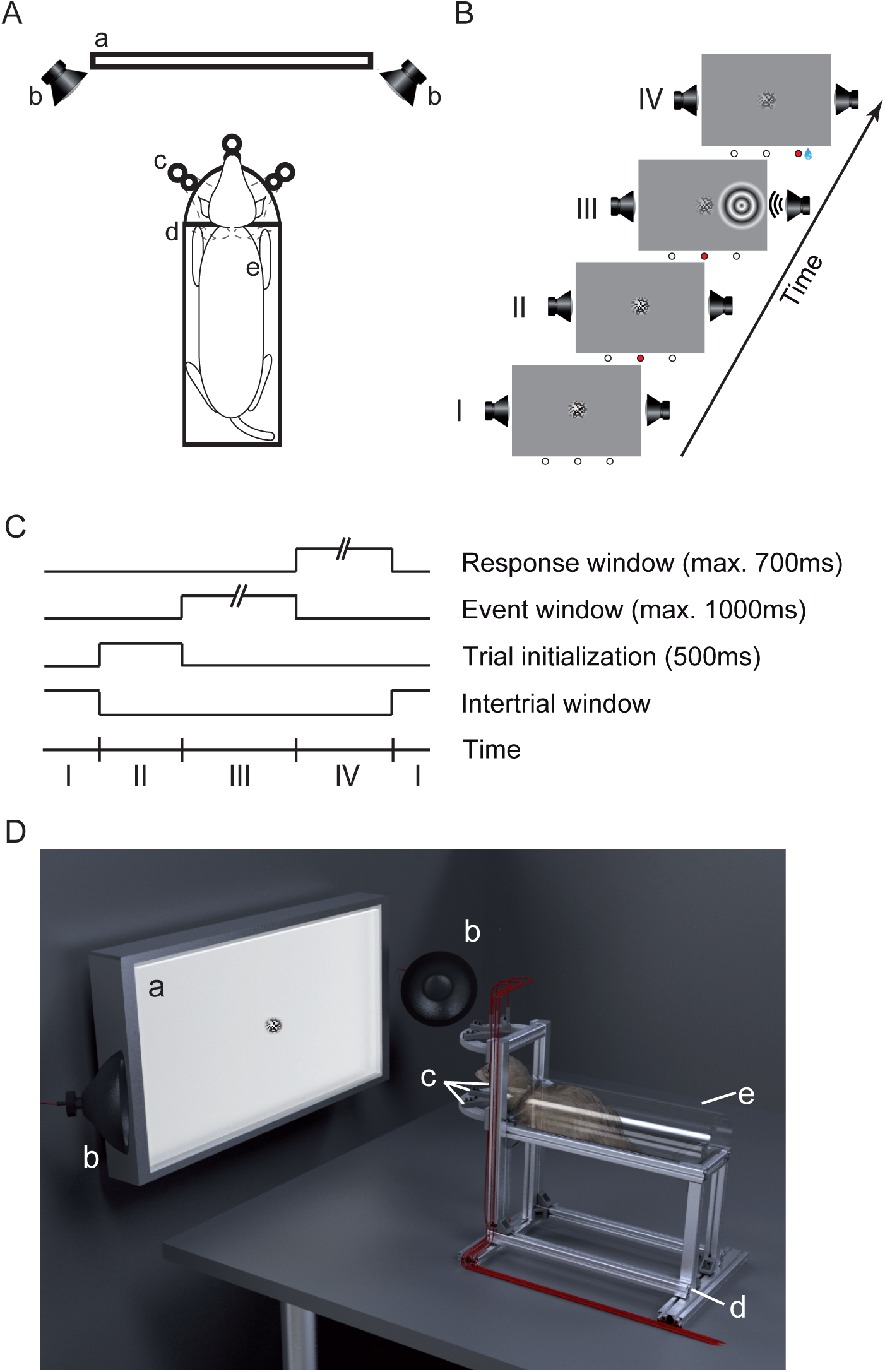
Experimental setup and behavioral task. (A) Schematic of the components of the experimental setup in a top view: the LED-screen (a) with a speaker (b) on each side, the aluminum pedestal (d), and the three light-barrier-waterspout combinations (c). The semi-circular acrylic tube with a ferret (e) inside was placed on the pedestal. (B) Successive phases in the detection task: The inter-trial window (I), the trial initialization window (II), the event window (III) and the response window (IV). The three circles below each frame represent the light-barriers (white = unbroken, red = broken). The center of the screen displays a static visual random noise pattern. (C) Schematic of trial timing. When the ferret broke the central light-barrier (II) for 500ms a trial was initialized and the event window started (III), indicated by a decrease in contrast of the static random noise pattern. At a random time between 0-1000ms during the event window the auditory and/or visual stimulus appeared for 100ms either left or right from the center. After stimulus offset the ferret had a response time window between +100-700ms (IV) to pan its head from the central position to the light-barrier on the side of the stimulation. Subsequently, the inter-trial screen (I) appeared again. During the whole session the screen’s global luminance remained unchanged. (D) Three-dimensional rendering of the experimental setup. Labeling of the components as in (A).

### Sensory stimulation

Auditory and visual stimuli were created using the Psychtoolbox (V3) [59] in a Matlab environment (The Mathworks Inc.; MA, USA). A white noise auditory stimulus (100ms) with up to 50dB sound pressure level (SPL) was used for auditory stimulation. It was generated digitally at 96kHz sample rate on a high-end PCI-audio card (HDSPe AES, RME-Audio, Germany) and delivered through two ‘T1’ Beyerdynamic speakers (Heilbronn, Germany). Visual stimuli consisted of concentric moving circular gratings (22.5°, 0.2cycles/°, 5Hz) up to 0.38 Michelson contrast (Cm) shown for 100ms (6 frames @ 60 Hz monitor-refresh rate). The background was set to half-maximum luminance to avoid global luminance changes at stimulus onset. In the center of the screen, a static random noise pattern was displayed (7°, Cm between 0 and 1). During ‘bimodal’ trials, both visual and auditory stimuli were presented synchronously as described below.

### Detection task

The ferrets were trained to solve a spatial detection task, as shown in Figure 1B and C. To initialize a trial the ferret had to maintain a central head position by breaking the central light-barrier for 500ms. This caused the random noise pattern in the center of the screen to decrease contrast and indicate to the animal that the stimulus-window (up to 1000ms) had started. During this interval the animal had to further maintain a central head position. A stimulus was presented for 100ms on the left or on the right side, respectively, starting at a random time in this window. The stimulus could be a unimodal visual (circular grating), unimodal audio (white noise burst) or temporally congruent bimodal stimulus (further details see below). After stimulus offset, the animal had to respond within 600ms by panning its head to the respective side. If the response was correct the animal received a water reward (~80μl) at that position and could immediately start the next trial. If the response was too early (before stimulus onset or within 100ms after stimulus onset), incorrect (wrong side) or omitted (no response), the trial was immediately terminated, followed by a 2000ms interval during which no new trial start was allowed.

### General procedure

Following the habituation to the harness, tube and setup all ferrets learned to detect unimodal stimuli. Two of the animals were trained in the auditory task first and then the visual; the other two were trained in reverse order. After completion of the training and reaching of sufficient performance, we presented stimuli of both modalities during the same sessions and determined the individual detection threshold. Twenty different stimulus amplitudes (0-50dB SPL; 0-0.38Cm) were chosen in a 1down/3up staircase procedure [60], i.e., if the animal solved the trial correctly (hits) the stimulus amplitude decreased by one step for the next trial, down to the minimum, whereas false responses (misses, or omitted responses) led to a 3 step increase. No change occurred for responses that were issued too early (rash trials). In each trial either the auditory or the visual stimulus was presented in a pseudo-randomized fashion with individual staircases. To avoid a side- or modality-bias, each modality-side-combination was titrated to an equal number of hits within each session. Due to the huge combinatorics of conditions, each ferret had to complete 10-15 sessions to accumulate a sufficient number of trials per amplitude level. The data of each animal were pooled and treated as one sample, i.e., session information was discounted during further analysis. Sensory thresholds were determined by fitting a Weibull function to the data for each ferret individually.

In a subsequent set of measurements, we combined simultaneous stimulus presentation in both modalities. To this end, we fixed the stimulus in one modality at the amplitude where the tested animal had an accuracy of 75% during the unimodal testing and varied the amplitude in the other modality according to the staircase procedure described above. In these bimodal sessions we again included the unimodal conditions, such that we obtained four different stimulation classes: unimodal auditory (A), unimodal visual (V), auditory supported by visual (Av), visual supported by auditory (Va). These four stimulation conditions were presented in a pseudo-randomized fashion and separate staircases during the sessions. All ferrets completed 10-12 sessions and the threshold was determined for each ferret by fitting a Weibull function to the data.

### Data Analysis

All offline data analysis was performed using custom written scripts in Matlab (The Mathworks Inc., MA, USA).

#### Psychometrics

We evaluated the accuracy values (P) for all N stimulus amplitude classes (a) with at least 6 hit trials in total on both sides using equation (1).

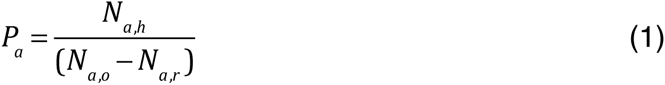

Here, *a* denotes the amplitude of the stimulus, *N_a,h_* (*hit trials*) was defined as the number of correct response trials for stimulus amplitude a, *N_a,o_* (*onset trials*) was the number of trials for stimulus amplitude a where the animal reached stimulus onset time, and *N_a,r_* (*rash trials*) as the number of trials for stimulus amplitude a were the animal gave a response before the response window had started (up to 100ms after stimulus onset), assuming the animal was guessing and not responding based on sufficiently collected sensory evidence. We estimated the detection threshold by fitting a Weibull function to P*_a_*,

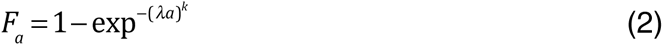

here *k* signifies the form-parameter and λ represents the scale-parameter. The number of trials were used as weights during the fitting procedure. Due to the fact that every animal had different thresholds in the respective modalities, we calculated the standard deviation of each fit by using a delete-d jackknife method, were d = 20% corresponds to the number of sessions excluded per run, i.e. 2 or 3, respectively.

#### Modeling cross-modal interaction

In order to quantify the cross-modal interaction, we used the MLE approach. Therefore we utilized the audio and visual accuracy from the multimodal experiment for all existing stimulus intensities. Assuming a model of a hidden Gaussian representation of the sensory input in the brain we estimated the variance (*σ*) for all points based on the *F_a_* values form the Weibull function,

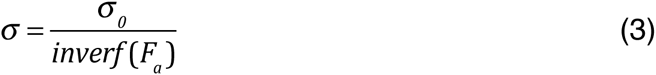

where ‘*inverf’* equates to the inverse error function and *σ*_0_ an unknown scale factor. As in the following calculation of *σ_bi_* it drops out we can set it arbitrarily to a value of 1. The next step was to combine both unimodal variances to derive the bimodal variance (*σ_bi_* ) according to

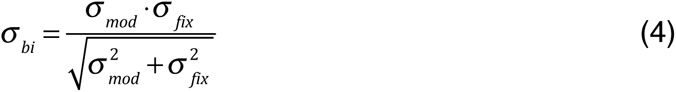

Where *σ_mod_* represents the variance for the modality which intensity were modulated and *σ_fix_* for the modality that was fixed at 75% threshold. Subsequently, we used the inverse value of the bimodal variance in an error function (*erf*) to determine the bimodal accuracy (5).

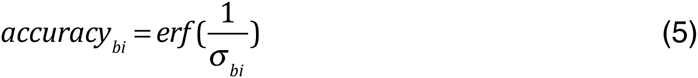

#### Reaction time

The RT was defined as the time difference between stimulus-onset and the time point when the animal panned its head out of the central light-barrier. Only intensity classes with at least 6 successful responses (hits) were included in the RT analyses. To quantify the RT differences between the corresponding amplitudes from uni- and bimodal stimulation we computed the Multisensory Response Enhancement (MRE) [49] as follows:
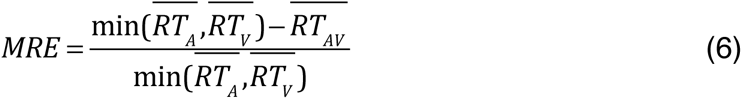

with 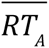 and 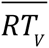 referring to the observed mean RT for the auditory and visual stimuli, respectively. 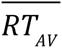 is the mean RT for the corresponding bimodal stimulus.

We calculated a race model [56] to evaluate potential RSE. In our study, accuracy varied across subjects and sensory conditions. In order to compare reaction times across subjects and compute the race model for all related modality combinations we introduced ‘subjective intensity classes’ (SIC) as determined by the accuracy fit in different unimodal conditions (0-74%, 75-89% and 90-100% indicating low, medium and high performance accuracy, respectively). This ensured a sufficient number of trials per SIC and additionally normalized for inter-individual differences in the range of stimulus amplitudes. Intensity and modality effects on the RT were tested applying the same grouping approach and computing a two-way ANOVA.

## Results

Four ferrets were trained in a lateralized audiovisual spatial detection task until they accomplished to solve the detection task in both modalities at high supra-threshold stimulus amplitudes (audio = 50dB SPL, visual = 0.38 Cm). The training was discontinued once the animals showed a stable baseline performance (>90%) across 5 consecutive days with high accuracy levels (audio = 92±1%, visual = 92±1%; mean±SEM). Two of the animals learned first the auditory (26 and 16 days training, respectively) and then the visual task (training for 28 days in both animals). The two other ferrets acquired the modalities in the opposite sequence (11 and 19 days for the visual and 14 and 14 days for the auditory modality, respectively). All animals achieved high performance levels demonstrating the viability of the training paradigm.

In all experiments for the determination of sensory thresholds we pooled results from left and right stimulation sides to calculate the accuracy values for all amplitudes. Testing for a laterality bias by comparing hit performance on both sides with a paired *t*-test revealed no significant bias (unimodal experiment: all animals = *p* >0.05; bimodal experiment: all animals = *p* >0.05).

### Determination of unimodal thresholds

In the first experiment we determined the 75% accuracy threshold for detection of visual and auditory stimuli in a unimodal setting for each individual ferret (Fig. 2), with an individual range of stimulus amplitudes for each animal. Ferrets performed on average 12 (±2) sessions (104±26 trials±SEM/session) in the unimodal experiment. Before pooling the sessions, we tested each ferret for non-stationarity effects across sessions by comparing the variance of the first three sessions at 84% accuracy threshold against the one of the last three sessions. We used three sessions as a minimum to ensure a sufficient number of trials for a proper Weibull function fit. No animal showed a non-stationarity in any modality (*p* >0.05 Two-sample t-test, 2-sided). The pooled data could well be described by a Weibull function (*r^2^* = 0.56 - 0.92, Fig. 2).

**Figure 2.**
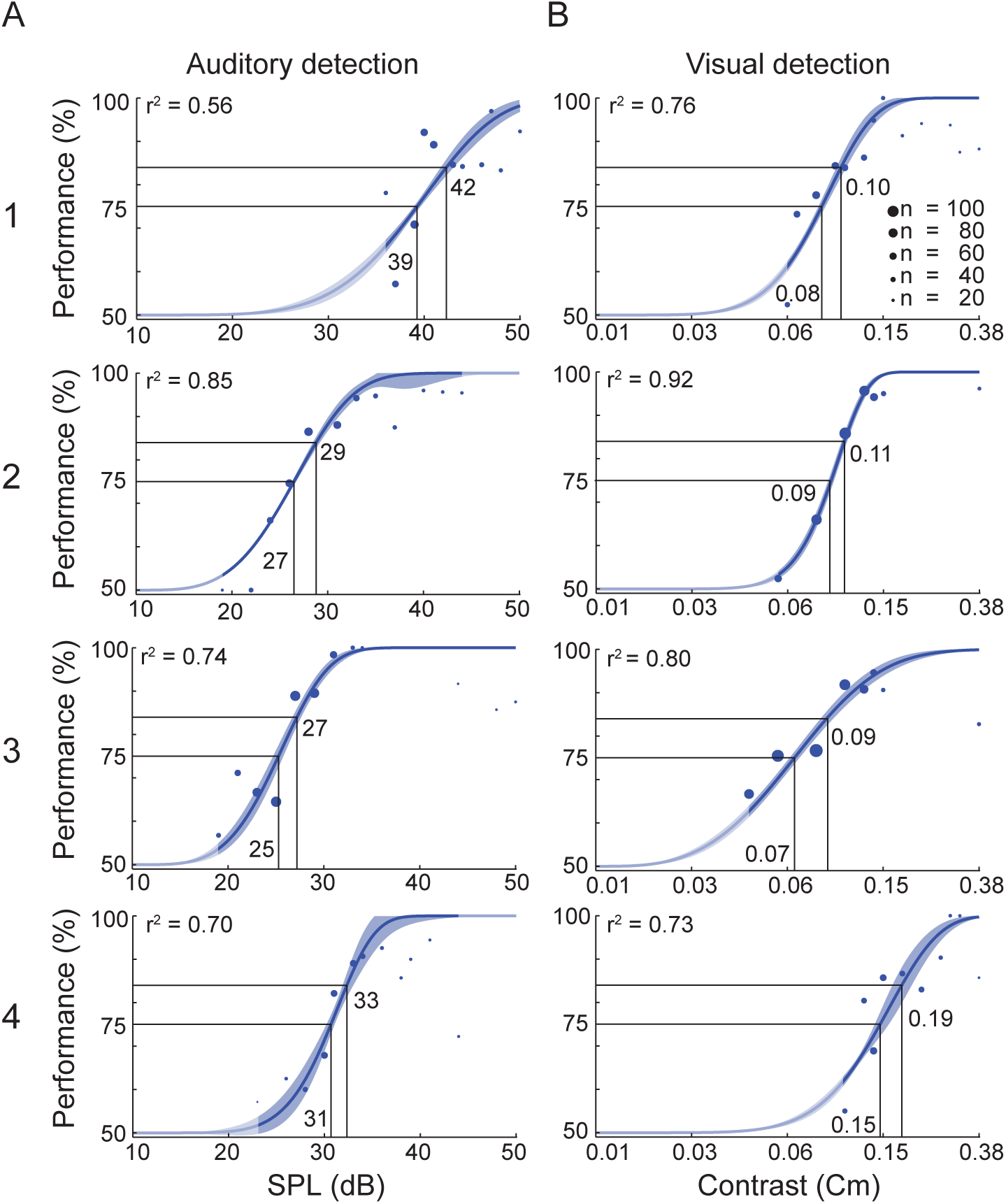
Detection task performance of the unimodal experiment. (A) Data for performance in the unimodal auditory detection task. (B) Data for the unimodal visual detection task. Each row represents one animal (1-4). Each dot represents the average performance of N trials (diameter) for the tested auditory amplitudes (dB SPL) or visual contrasts (Cm). The data are fitted by a Weibull function. Numbers within the panels indicate the amplitude values corresponding to the 75% and 84% thresholds, respectively. The blue shaded area around the fit indicates the standard deviation. The unmasked parts of the graphs indicate the range of the actually tested stimulus amplitudes.

### Determination of uni- and bimodal thresholds

In the second experiment, the two crossmodal stimulation conditions were added to the sessions. One modality’s intensity was fixed at 75% threshold, as determined from the unimodal experiment (Fig. 2) while the other modality was varied in amplitude according to a staircase procedure. All ferrets participated in 12 (±1) multimodal sessions (111±37 trials±SEM/session). Like for the unimodal sessions, we again tested for non-stationarity effects between the first and the last sessions by comparing the 84% accuracy threshold variance as determined by the Weibull fit. Since the introduction of bimodal classes reduced the relative number of unimodal stimulus presentations during each session, we had to pool minimum across the first and last 5 sessions, respectively, to generate a proper Weibull fit. No animal showed non-stationarity across the bimodal sessions (2-sided two-sample *t*-test; *p*>0.05). Subsequently, we calculated the accuracy for each amplitude where at least 6 trials had been performed and the psychometric curves were fit using a Weibull function (Fig. 3). The pooled data could well be described by a Weibull function (*r^2^* = 0.39 - 0.90, Fig. 3).

**Figure 3.**
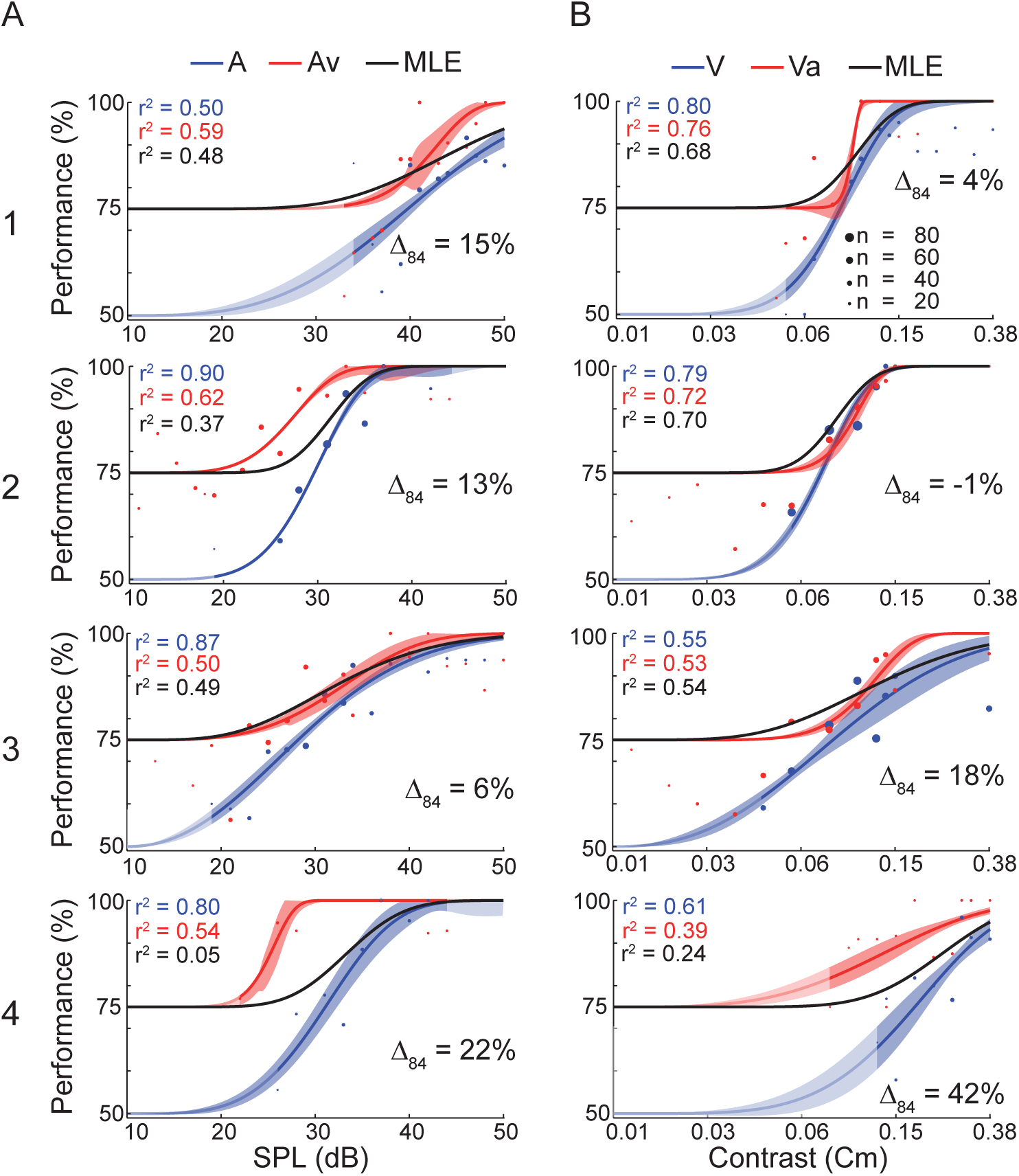
Detection task performance of the bimodal experiment. (A) Data for the stimulus conditions auditory-only (A) and auditory stimulation supported by a visual stimulus (Av). (B) Data for the stimulus conditions visual-only (V) and visual stimulation supported by an auditory stimulus (Va). Each row represents one ferret (1-4). Each dot represents the average performance of N trials (diameter) at a given auditory amplitude (dB SPL) or visual contrast (Cm). The data are fitted by a Weibull function. The uni- and bimodal fit is represented by the blue and red line, respectively. The shaded area around the fit indicates the standard deviation. Δ_84_ displays the relative amount of threshold shift of the bimodal compared to the unimodal psychometric function at a performance of 84%. A positive shift indicates a threshold decrease. The black curve represents the MLE model. The unmasked parts of the graphs indicate the range of the actually tested stimulus amplitudes.

The comparison of the unimodal 75% thresholds between both experiments revealed a slight increase from the uni- to the multimodal experiment, except in animal 2 which showed a decrease (Table 1). However, the differences were smaller than one of the respective amplitude steps in the staircase procedure. Furthermore, two of the animals (1 and 4) did not reach a performance above 90±5% in the highest intensities in one modality (audio and visual, respectively). These findings indicate that the bimodal experiments were slightly more demanding, presumably because four stimulation conditions were presented compared to the unimodal experiment with only two stimulation conditions.

**Table 1.** Comparison of threshold values for uni- and bimodal experiments.

|  | Amplitude values @ 75% |  | Amplitude values @ 84% |  |  |
| --- | --- | --- | --- | --- | --- |
|  | unimodal Exp. | bimodal Exp. | unimodal Exp. | bimodal Exp. |  |
|  | A | A | A | A | Av |
| 1 | 39 | 40 | 42 | 45 | 41 |
| 2 | 27 | 30 | 29 | 32 | 26 |
| 3 | 25 | 28 | 27 | 33 | 31 |
| 4 | 31 | 31 | 33 | 34 | 25 |
|  | V | V | V | V | Va |
| 1 | 0.08 | 0.08 | 0.10 | 0.10 | 0.09 |
| 2 | 0.09 | 0.08 | 0.11 | 0.09 | 0.09 |
| 3 | 0.07 | 0.10 | 0.09 | 0.18 | 0.12 |
| 4 | 0.15 | 0.19 | 0.19 | 0.25 | 0.10 |

The amplitude values at the 75% and 84% thresholds (in dB SPL for A and Av; Cm for V and Va) in the unimodal and bimodal experiments (columns) for all animals (rows 1-4).

Because different values were used for the lower bounds in uni- (50%) and crossmodal (75%) fitting, we employed the 84% threshold for comparison of performance between uni-and crossmodal settings. All fits to the bimodal psychometric functions showed a left shift compared to their unimodal complements, except for animal 2 in the V-Va comparison (amplitude decrease ±SEM: A-Av = 5.3±1.5; V-Va = 0.06±0.03; for absolute values see Table 1). This demonstrates a decrease in detection thresholds in all ferrets, except for animal 2 in the Va condition where the auditory stimulus had no augmenting effect. For quantification we calculated the relative shifts at the 84% performance-level between the uni- and bimodal psychometric fit (Δ_84_ in Fig. 3). A positive number indicates a lower threshold as determined by the bimodal fit, i.e., an increase in bimodal detection performance. On average, there was a shift (±SEM) of 15±5%, indicating an effective bimodal integration.

### Maximum likelihood estimates

To investigate whether ferrets integrate the two sensory modalities in a statistically optimal fashion, we computed a MLE model and compared the *r^2^*-difference between the empirical data (Fig. 3, red) and model (Fig. 3, black). The range of the difference Δ*_bimodal–MLE_* was -1 to49% (mean difference ±SEM 14±6). In four cases the MLE matched the bimodal psychometric function and the difference of the explained variance between the empirical fit(Fig. 3) and the MLE was 10% or less (A1: Δ*_Va–MLE_* = 8%; A2:Δ*_Va–MLE_* = 2%; A3:Δ*_Av–MLE_* = 1% andΔ*_Va–MLE_* = -1%). For one condition ( animal 1:Δ*_Av-MLE_*= 11%) the MLE underestimated the empirical fit at the highest stimulus amplitudes (Fig. 3A). This may be caused by the low unimodal performance at high stimulus amplitudes, since the MLE model depends on the unimodal performance. This argument also holds true for the Va case (Δ*_Va–MLE_* = 15%) of animal 4 (Fig. 3B, bottom panel). If the animal had shown a unimodal performance comparable to that previously measured in the unimodal experiment, the MLE model would be similar to the empirical bimodal fit. In the other two cases the MLE underestimated the empirical fit in the intermediate amplitude ranges (animal 2: Δ*_Av–MLE_* = 25% and animal 4: =49%, Fig. 3). Overall, the MLE modeling results support the conclusions drawn from the comparison of the 84% performance threshold between uni- and bimodal conditions. The results indicate that ferrets integrated the two modalities as good or even better than predicted by the MLE estimator (Fig. 3).

### Reaction time analysis

One of the most important benefits of multisensory integration is the reduction of RTs for bimodal stimuli compared to unimodal stimulation. The measured RT varied during the multisensory experiment with target amplitude in all modality types. In all stimulus conditions and all animals, RT showed a significant negative correlation with stimulus amplitude (range A: *r* = -0.17 to -0.41; V: *r* = -0.25 to -0.45; Av: *r* = -0.21 to -0.44; Va: *r* = -0.34 to -0.46; all correlations: *p* < 0.01; Fig. 4). RT significantly increased with decreasing amplitude (ANOVA *p* < 0.05) in all but one condition (animal 1: audio-alone, ANOVA *p* > 0.05). This is an expected finding, because the signal-to-noise ratio (SNR) decreases with decreasing stimulus amplitude and the internal signal processing is slower for low SNR.

**Figure 4.**
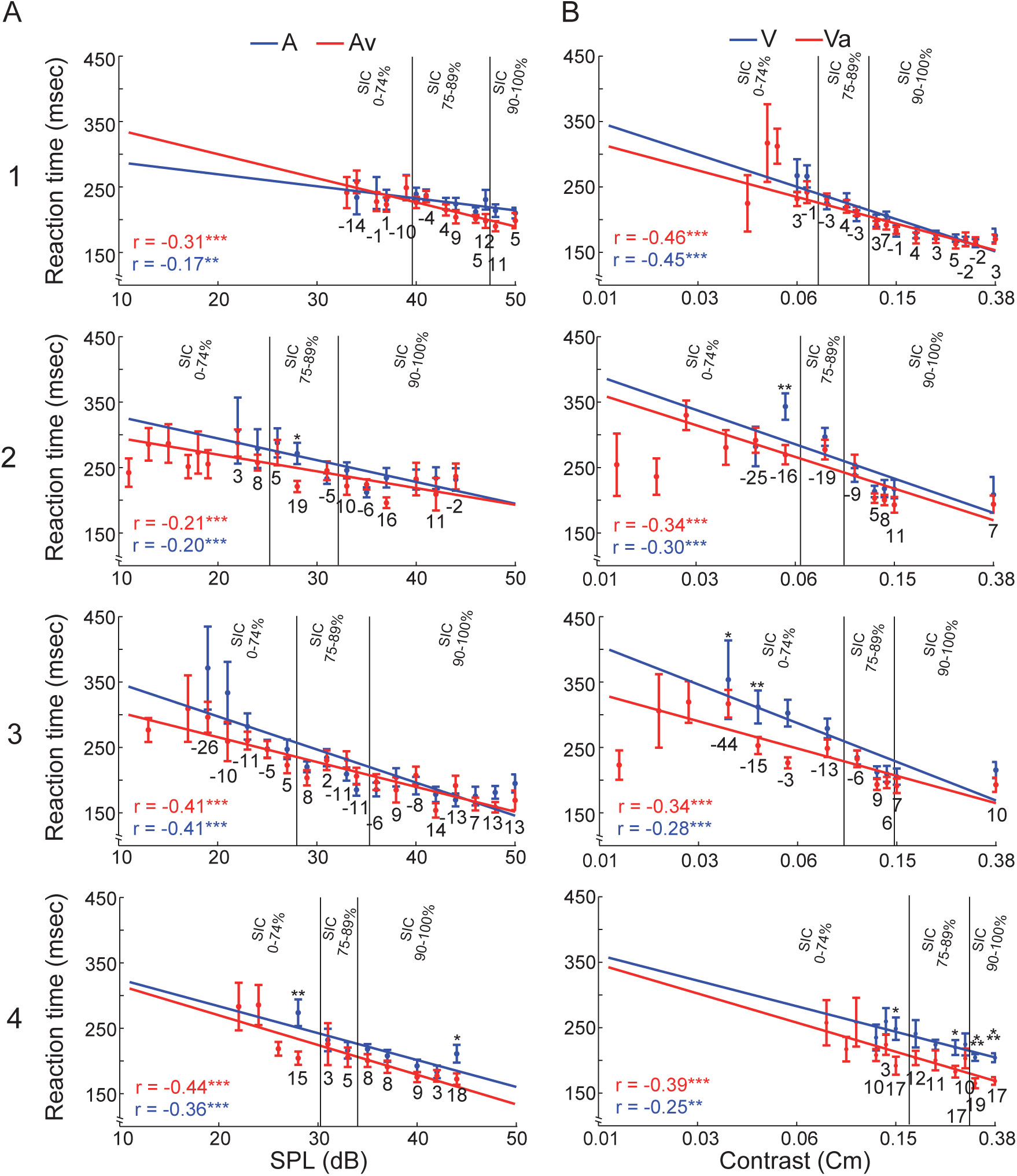
Reaction time data from the bimodal experiment. (A) Data for the stimulus conditions auditory-only (A) and auditory stimulation supported by a visual stimulus (Av). (B) Data for the stimulus conditions visual-only (V) and visual stimulation supported by an auditory stimulus (Va). Each row represents one ferret (1-4). RT ± SEM are shown as a function of stimulus amplitude (red = bimodal, blue = unimodal). Each data point represents the RT average for all hit trials recorded at that amplitude. Asterisks indicate significant differences between uni- and bimodal conditions (*t*-test: * = *p* < 0.05, ** = *p* < 0.01, *** = *p* < 0.001). Below each pair of uni- and bimodal RTs the Multisensory Response Enhancement (MRE) is shown as numerical values. In each panel, Pearson correlation coefficient and regression line for both data sets are shown. The two vertical lines mark the borders between the subject intensity classes (left of first line: 0-74%, between the lines 75-89%; right of the second line 90-100% performance).

To reduce the dimensionality and compare reaction times across subjects we used ‘subjective intensity classes’ (SIC) (see Material and Methods). To quantify RT changes reflecting potential multimodal enhancement effects, we calculated the MRE for all uni- and bimodal stimulus pairs and summed these according to the SICs. The average MRE of both modalities was slightly positive (Av = 3.59%; Va = 0.06%). However, about one-third of the cases (7 out of 24, Table 2) showed a negative MRE. Such negative MRE values, which indicate that the average unimodal RT is faster than the average RT of the bimodal condition, occurred only in the low and medium SIC . In the highest SIC, the MRE was consistently positive. Overall, the MRE results suggest a multimodal enhancement effect in the high and medium and an interfering effect in the lower SIC.

**Table 2.** Reaction time: average MRE.

|  | 0-74% | 75-89% | 90-100% | 0-74% | 75-89% | 90-100% |
| --- | --- | --- | --- | --- | --- | --- |
| | MRE $A_v$ | | | MRE $V_a$ | | |
| 1 | -6.00 | 4.33 | 8.00 | 1.00 | -0.67 | 2.22 |
| 2 | 5.50 | 6.33 | 5.50 | -20.50 | -19.00 | 4.40 |
| 3 | -9.40 | -3.00 | 3.63 | -18.75 | 3.00 | 8.50 |
| 4 | 15.00 | 4.00 | 9.20 | 10.00 | 12.50 | 18.00 |

Multisensory Response Enhancement (MRE) computed for the RTs from all animals (rows) and stimulus conditions of the bimodal experiment according to equation 6. (see Methods). The MRE’s were sorted by the subjective intensity classes (SIC ; columns from left to right). Av: auditory supported by visual; Va: visual supported by auditory.

To investigate a potential RSE we calculated a race model on the pooled RTs according to the SICs. The race model assumes that during multimodal stimulation no modality integration happens, but that signals of either modality are processed independently. Whichever of the two leads to a result first triggers and determines the response, i.e., the head movement towards the detected stimulus. Therefore , the bimodal cumulative distribution function (CDF ) of the RT can be modeled by sampling from the unimodal RT CDFs. Afterwards the modeled bimodal RT CDF can be compared with the empirical bimodal RT CDF (see Fig. 5). If the empirical RT CDF is faster in 20-50% of the percentiles compare to the modeled RT CDF the race model can be rejected and modality integration is suggested [61]. For a detailed explanation of the race model see Ulrich et al. [56].

**Figure 5.**
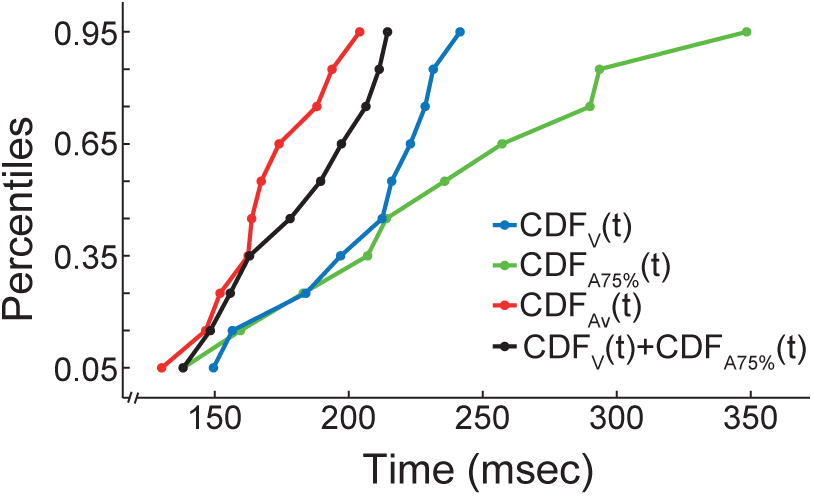
Race model example. Analysis of RT CDFs from animal 4. High visual SIC CDFs are shown for unimodal visual stimulation (V, blue), auditory stimulation at 75% (A75%, green), auditory stimulus supported by visual stimulation (Av, red) and the combination of both unimodal CDFs (V+A75%, black). In this case the race model gets rejected, because the empirical bimodal CDF (red) is ‘faster’ than the modeled CDF (black).

We computed the relative (%) deviation from the linear unimodal combination for all stimulus conditions (Fig. 6) for each SIC. If this difference for the empirical bimodal CDF is in 20-50% of the cases negative the race model can be rejected (Miller and Ulrich, 2003). The biggest effect of the supportive value occurred in the highest intensity group, because there the change was negative compare to the combined unisensory CDF in the lower percentiles (upper row, Fig. 6). In the 75-89% SIC no percentile of the crossmodal combinations was negative (middle row Fig. 6) and in the lowest intensity-group the bimodal and the supportive value RTs were similar (bottom row Fig. 6), i.e., the benefit of the redundant signal seems to diminish with decreasing intensity group. However, in the medium and high performance classes the bimodal RT seemed to be closer to the combined CDF than each of the unimodal distributions. For the high SICs, the distributions suggest that the race model can be rejected at a descriptive level. Overall, these results are compatible with the notion that, for higher SICs, multisensory integration processes are leading to RT gains beyond what can be predicted from the fastest unimodal responses.

**Figure 6.**
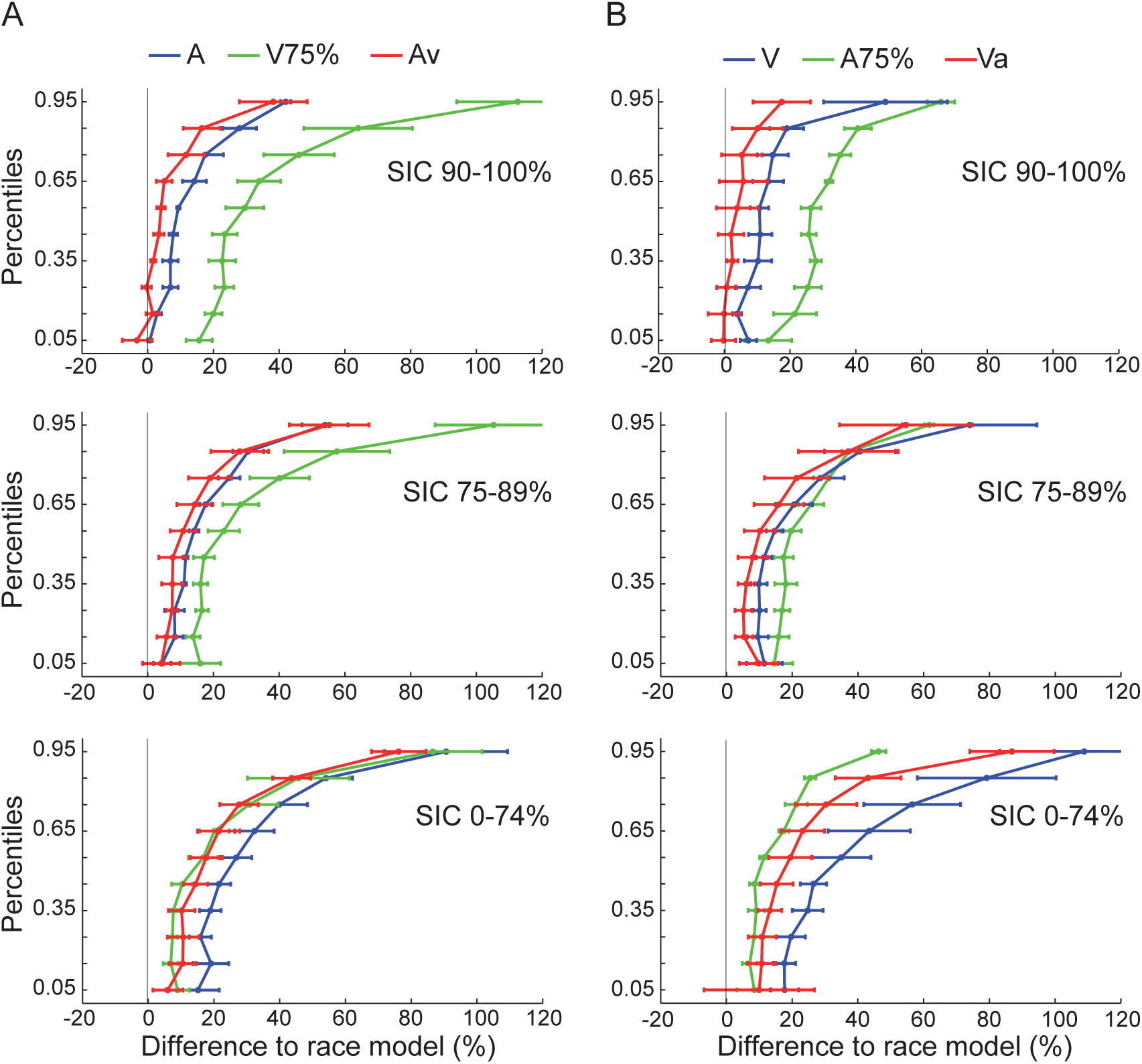
Reaction time: race model results. The RTs were sorted by the SICs (rows) and both modalities (A: audio, B: visual) pooled across all animals. The X-axis displays the cumulative reaction time differences to the race model for each modality (± SEM). A value of 0 at the X-axis corresponds to the prediction from the combination of both unimodal CDF’s. The blue curve displays the unimodal condition, the green curve the RTs at the supportive value and the red curve the bimodal class, respectively.

To investigate intensity, modality and interaction effects on a more global scale we pooled the RT of all animals according to subjective intensity classes and calculated a two-way ANOVA, with modality and intensity as main factors (Fig. 7). This revealed main effects in both factors (Modality: *F*(3, 4632) = 18.84 (*p* < 0.001); Intensity: *F*(2, 4633) = 310.65 (*p* < 0.001)) and an interaction effect (Modality* Intensity: *F*(6, 4624) = 3.93 (*p* < 0.01)). A post hoc *t*-test (Holm-Bonferroni corrected) revealed significant differences between and within performance classes (Fig. 7), respectively. The post hoc *t*-tests between the intensity groups and modalities were all highly significant (*p* < 0.001). This result suggests that the ferrets’ RTs increase as the intensity of the stimulus gets weaker and significantly decrease in the multimodal compared to the unimodal classes.

**Figure 7.**
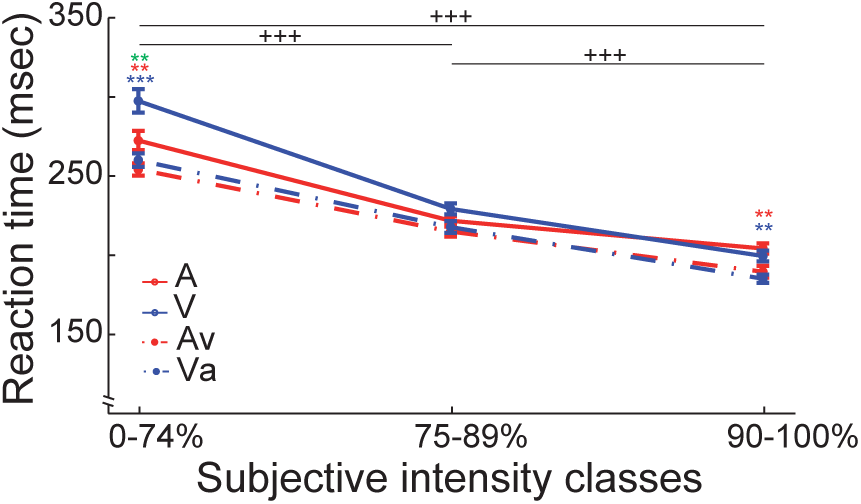
Reaction time: two-way ANOVA results. The reaction times (RT) pooled by subjective intensity classes (0-74%, 75-89%, 90-100%). The X-axis displays the three performance classes and the Y-axis shows the RT in milliseconds ± SEM. The solid lines represent the unimodal, the dashed lines the bimodal, red indicates the audio and blue the visual modalities (*: *p* < 0.05; **: *p* < 0.01; ***: *p* < 0.001; Holm-Bonferroni corrected). +++, significant differences between performance classes within each modality (Holm-Bonferroni corrected); red and blue asterisks, significant differences between uni- and bimodal conditions in one performance class (Holm-Bonferroni corrected); green asterisk, significant difference between the two unimodal conditions.

## Discussion

How information from different modalities is integrated has been a subject of intense research for many years. Here we asked if ferrets integrate sensory signals according to the same principles established for other species [31,33,35–39,47,62,63]. We expected the ferrets to perform more accurately and with lower RTs in the bimodal cases, because congruent inputs from two modalities provide more reliable sensory evidence [62,64–66]. As predicted, bimodal detection thresholds were reduced and RTs were faster in the bimodal compared to unimodal conditions, demonstrating multimodal integration effects. Furthermore our results on MLE modeling suggest that ferrets integrate modalities in a statistically optimal fashion.

### Methodological considerations

Previous studies in behaving ferrets have used either freely-moving [1,13,15,55] or head-restrained [26] animals. Here, we developed a head-free, body-restrained approach allowing a standardized stimulation position and the utilization of the ferret’s natural response behavior. The setup is especially suited for psychometric investigations because the distance between animal and the stimulus sources remains constant across trials. The high inter-trial-consistency and the fixed animal position allow the combination of behavioral protocols with neurophysiological recordings comparable to head-restrained approaches [26]. An additional advantage is the usage of a screen instead of a single light-source for the visual stimulation [1,31], enabling the spatially flexible presentation of a broad variety of visual stimuli. Similar to other ferret studies [13,55], one limitation of our approach lies in the relatively low number of trials collected per session. We therefore had to pool data from different sessions to obtain a sufficient number of trials for the fitting of psychometric functions. Merging of sessions was justified by the absence of non-stationarity effects and the high amount of variance explained by the fits. This also indicates a low day-to-day variability of perceptual thresholds. Our results complement that of an earlier study in ferrets demonstrating that measured thresholds were not affected by trial-to-trial fluctuations in the animals’ decision criterion [1]. Overall, these findings suggest that the experimental design presented in this study is well suited for psychophysical investigations.

Establishing links across species, our behavioral paradigm was inspired by previous human psychophysical studies which showed that temporally congruent crossmodal stimuli enhance detection [62,64–66]. Frassinetti et al. [62] adopted an animal approach [51] to humans and obtained similar results in terms of multisensory enhancement effects. Another study form Lippert and colleagues [64] showed that informative congruent sounds improve detection rates, but this gain disappears when subjects are not aware of the fact that the additional sound offers information about the visual stimulus. They concluded that cross-modal influences in simple detection tasks are not exclusively reflecting hard-wired sensory integration mechanisms but, rather, point to a prominent role for cognitive and contextual effects. This contrasts with more classical views suggesting that information form different sensory modalities may be integrated pre-attentively and substantially rely on automatic bottom-up processing [35]. Our observation of the inter-experiment threshold increase for the unimodal conditions might suggest possible contextual effects. A possibility is that, in the second experiment, the inclusion of the bimodal conditions may have created a contextual, or motivational, bias of the animals towards solving the bimodal trials because more sensory evidence was provided. This could also explain why the performance in the unimodal conditions of the bimodal experiment did not reach 95-100% accuracy even at the highest intensities, unlike in the unimodal experiment.

Taken together, our study demonstrates that the implemented behavioral paradigm is suitable to determine uni- and bimodal thresholds and to operationalize multisensory integration processes. Possible contextual and attention-like effects seem hard to elucidate by pure psychometrics, but simultaneous electrophysiological recordings could provide valuable insights into the underlying brain processes during the task.

### Optimal modality integration

This is the first study on behaving ferrets to quantify multimodal enhancement effects and to test for optimal modality integration. The results of our bimodal experiment show clear multisensory enhancement effects. The left shift of the psychometric function and the variance reduction, derived at 84% accuracy, demonstrate increased detection rates and enhanced reliability for lower test-intensities in the bimodal stimulation conditions, indicating that the ferrets indeed integrate information across modalities as shown for other species [31,35,37,47,54,63–65]. MLE modeling is typically used in multisensory integration to test the hypothesis that the integrative process is statistically optimal by fitting the parameters of the model to unisensory response distributions and then comparing the multimodal prediction of the model to the empirical data. Studies on humans have shown that different modalities get integrated in a statistical optimal fashion. For example, Battaglia et al. [37] found that human subjects integrate audio and visual modalities as an optimal observer. The same is true for visual and haptic integration [36], and integration of stereo and texture information [39,67]. Furthermore, Alais and Burr [38] could show that the ventriloquist effect is based on near-optimal sensory integration. Rowland and colleagues showed statistical optimal integration in the cat for audio-visual perception [63] and Gu et al. [47] could demonstrate the same principle in macaques for visual and vestibular sensory integration. Similar to the abovementioned studies, our results on MLE modeling suggest that ferrets integrate modalities in a statistically optimal fashion. Surprisingly, in two of our cases the MLE underestimates the empirical fit, which is counterintuitive because the MLE provides already the maximum estimate. A potential explanation might be that multisensory benefit is larger for some modalities compared to others, as suggested by the modality precision hypothesis by Welch and Warren [68]. These hypotheses states that discrepancies are always resolved in favor of the more precise modality, i.e. the modality with the highest SNR gets weighted higher in the final sensory estimate. Battaglia and coworkers [37] showed that humans have a bias towards the visual modality in a multisensory spatial detection task. Finally, it could be caused by a low unimodal performance in the intermediate intensities since the MLE model depends on the unimodal performance. In summary, the MLE model provides evidence that ferrets merge modalities in a near-optimal fashion, similar to other species [36–38,47,67].

### Multisensory response enhancement

In a second analysis approach we compared RTs of the uni- and bimodal stimulation conditions and computed a race model to test a RSE. Our main results are in line with findings from other species. Previous work in humans revealed that subjects respond faster to bimodal compared to unimodal stimuli [49,64]. Miller [53] showed that this RT gain is a result of a modality integration effect and not only a product of the fastest processed modality. Gleiss and Kayser [31] demonstrated that additional non-informative white noise decreases RT in a visual detection task in rats. The effect size of the RT gain increased when the light intensity decreased. In our study the influence of amplitude on RT is directly related to the SNR, i.e., the internal signal processing is faster for high SNR. For high intensities of the varying modality (>75% unimodal performance), the SNR should be higher compared to the fixed supporting modality. Decreasing the intensity of the variable modality leads to a continuous decrease of its SNR (until 0), such that for low intensities the RT is completely determined by the amplitude of the supporting modality. Interestingly, some MRE values were negative in the lower and intermediate subjective intensity classes. This is due to the fact that the MRE model uses the fastest unimodal RT for calculation and the RT of the supporting values is faster than the average bimodal RT. The variable modality seems to have a competitive effect on the RT at low intensities, because the average bimodal RT is slower than the RT of the supportive value.

In addition to the MRE analysis, we computed a race model for the RT data. The race model tests RT effects in a more sophisticated way, by comparing a modeled bimodal RT CDF with the empirical bimodal RT CDF. In our dataset, the benefit of the redundant signal increased from low to high SIC. Data reached the criterion to reject the race model only in the high SIC. In the intermediate and low SIC the linear unimodal combination was faster compared to the empirical bimodal conditions. Nevertheless, in the intermediate SIC the bimodal percentiles were closer to the linear combination than the unimodal groups, indicating a minor gain of the supportive value and therefore a multisensory enhancement effect. In contrast, in the low SIC the bimodal group matches the supporting value group, implying that the supportive value is the driving modality in the sensory process [57,61] .

### Conclusions

In conclusion, our data demonstrate that basic principles of multisensory integration, such as enhancement effects of bimodal stimuli on detection rates, precision and RT apply to crossmodal processing in the ferret brain. The race model and MLE modeling provide evidence that ferrets integrate modalities in a statistically optimal fashion. To quantify this in more detail more advanced behavioral paradigms would be required where the stimulus onset varies across modalities and a broader range of stimulus amplitudes of supporting modality can be covered.

The setup we have developed to test ferrets in uni- and bimodal conditions is similar to human and non-human primate tasks, and can be combined in future research with approaches for the study of the underlying neural processes. Our behavioral paradigm could be combined with neuroscientific approaches such as, e.g., optogenetics or *in vivo* imaging [69]. Furthermore, the same setup could be used to implement more complex behavioral paradigms such as discrimination or go/no-go tasks [26]. Moreover, the setup would also be suitable to investigate aspects of sensory processing other than multisensory integration relating, e.g., to altered developmental conditions [7,12,24], to top-down influences on sensory processing, or to large-scale communication across distinct sensory regions during different behavioral paradigms. Altogether, our results describe a highly multifunctional experimental approach, which may further enhance the viability and suitability of the ferret model.

## Acknowledgements

We would like to thank Dorrit Bystron for assistance during data acquisition and Guido Nolte for advice in data modeling.

## Supporting Information

**S1 ARRIVE Guidelines.** Completed “ARRIVE Guidelines Checklist” for reporting animal data in this manuscript.

**S2 Dataset. Raw data of the unimodal detection task for all animals.**

**S3 Dataset. Raw data of the multimodal detection task for all animals.**

